# Digital data repository and automatic analysis framework for FDOPA PET neuroimaging

**DOI:** 10.1101/2022.04.14.488129

**Authors:** Giovanna Nordio, Rubaida Easmin, Alessio Giacomel, Ottavia Dipasquale, Daniel Martins, Steven Williams, Federico Turkheimer, Oliver Howes, Mattia Veronese

## Abstract

**Introduction:** FDOPA PET has been used extensively to image the human brain in many clinical disorders and has the potential to be used for patient stratification and individualized treatment. However, to reach its full and effective clinical translation, FDOPA PET requires both a robust data infrastructure and analytical protocol that are capable of ensuring high quality data and metadata, accurate biological quantification, and replicable results. In this study we evaluate a digital data repository and automated analytical framework for FDOPA PET neuroimaging that can produce an individualised quantification of dopamine synthesis capacity in the living human brain.

**Methods:** The imaging platform XNAT was used to store the King’s College London institutional brain FDOPA PET imaging archive, alongside individual demographics and clinical information. A fully automated analysis pipeline for imaging processing and data quantification was developed in Python and integrated in XNAT using the Docker technology. *Reproducibility* was assessed in test-retest datasets both in controls and patients with psychosis. The agreement between the automated analysis estimates and the results derived by the manual analysis were compared. Finally, using a sample of healthy controls (N=115), a *sensitivity* analysis was performed to explore the impact of experimental and demographic variables on the FDOPA PET measures.

**Results:** The final data repository includes 892 FDOPA PET scans organized from 23 different studies, collected at five different imaging sites. After removing commercials studies, the infrastructure consisted of 792 FDOPA PET scans from 666 individuals (female 33.9%, healthy controls 29.1%) collected from four different imaging sites between 2004-2021. The automated analysis pipeline provided results that were in agreement with the results from the manual analysis, with a Pearson’s correlation that ranged from 0.64 to 0.99 for Ki^cer^, and from 0.79 to 1.00 for SUVR. The mean absolute difference between the two pipelines ranges from 3.4% to 9.4% for Ki^cer^, and from 2.5% to 12.4% for SUVR. Moreover, we found good reproducibility of the data analysis by the automated pipeline (in the whole striatum for the Ki^cer^: ICC for the controls = 0.71, ICC for the psychotic patients = 0.88). From the demographic and experimental variables assessed, gender was found to most influence striatal dopamine synthesis capacity (*F = 10.7, p <0.001*), with women showing greater dopamine synthesis capacity than men, while the effects of weight, age, injected radioactivity, and scanner, varied by brain region and parameter of interest.

**Conclusions:** Combining information from different neuroimaging studies has allowed us to test comprehensively the automated pipeline for quantification of dopamine synthesis capacity using FDOPA PET data and to validate its replicability and reproducibility performances on a large sample size. This validation process is a necessary methodological step for the development of the clinical application of FDOPA PET as precision medicine biomarker. The proposed infrastructure is generalisable behind the FDOPA radiotracer.

## INTRODUCTION

Positron emission tomography (PET), in combination with the 6-[18F]fluoro-L-dopa (FDOPA) radiolabelled tracer, has been extensively used to image the dopamine system in vivo in living human brain (Youdim, Edmondson, and Tipton 2006). Accumulation of FDOPA in the brain parenchyma reflects its transport, decarboxylation into labelled dopamine, and vesicular uptake in the nigrostriatal presynaptic nerve terminals (**Figure 1A**). The tracer was introduced in 1983 (Garnett, Firnau, and Nahmias 1983) to quantify the integrity of the nigrostriatal dopamine and it found immediate application in subclinical models of dopamine neuronal damage and in Parkinson’s Disease (PD) studies (Calne et al. 1985; Nahmias et al. 1985). However, it took 39 years before it received FDA approval as an imaging agent to visualize dopaminergic nerve terminals in the striatum of patients with suspected Parkinsonian syndromes (Center for drug evaluation and research n.d.). Besides PD and dopaminergic neurodegeneration, FDOPA PET has proved to be valuable in other medical specialities. In neuro-oncology FDOPA PET has been used as an amino-acid tracer to detect both primary and recurrent gliomas, outperforming standard PET with 2-deoxy-2-[fluorine-18]fluoro-D-glucose integrated with computed tomography (18F-FDG PET/CT) in terms of both accuracy and sensitivity for classification between high-grade from low-grade gliomas (Somme et al. 2020; Xiao et al. 2019). In psychiatry, FDOPA PET has been extensively used to quantify the dopamine system in the pathophysiology of psychotic and other symptoms across conditions, including schizophrenia (Davis et al. 1991; Howes and Kapur 2009), bipolar disorder (Jauhar et al. 2017), 22q11 syndrome (Rogdaki et al. 2021), attention deficit disorders (ADHD) (Ernst et al. 1998), and substance dependence (Bloomfield et al. 2014). Several lines of evidence have linked FDOPA PET to treatment response in psychosis (Avram et al. 2019; Howes et al. 2012; Jauhar, Veronese, Nour, Rogdaki, Hathway, Turkheimer, et al. 2019; Kim et al. 2017) suggesting that it might be used as a neurochemical basis to discriminate between patients likely to respond and those unlikely to respond to first-line antipsychotic drugs (Veronese, Santangelo, et al. 2021).

**Figure 1:**
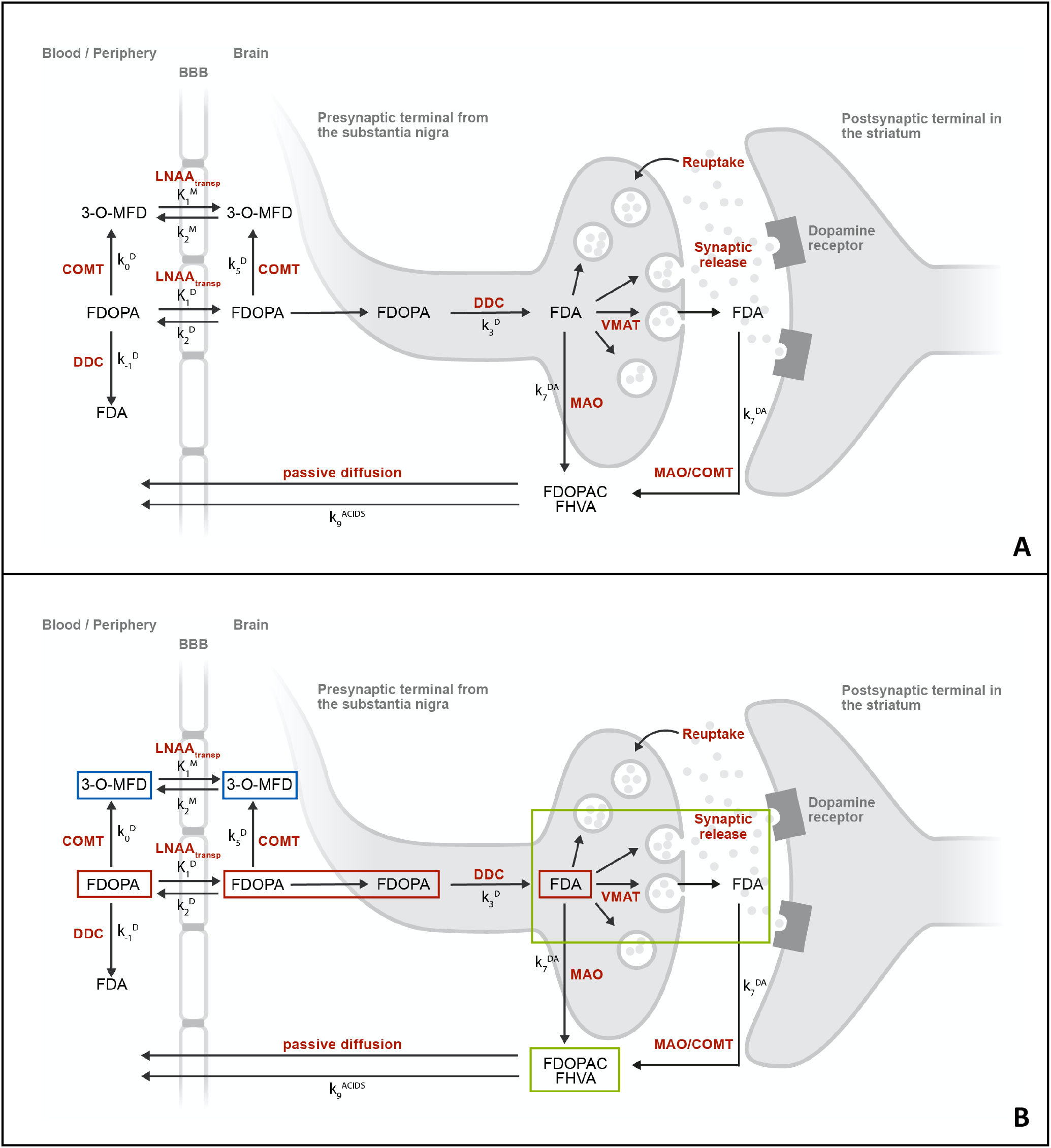
**A) [^18^F]Fluorodopa (FDOPA) PET tracer kinetics in brain and periphery.** After intravenous injection, the FDOPA in circulation is O-methylated at apparent rate constant k_0_^D^ (min^-1^) by cathecol-O-methyltransferase (COMT) to form 3-O-methyl-fluorodopa (3-O-MFD). Alternatively, the FDOPA in circulation can be decarboxylated at apparent rate constant k_-1_^D^ (min^-1^) by the enzyme DOPA decarboxylase (DDC) to form [^18^F]fluorodopamine (FDA). Both FDOPA and its COMT metabolite are subsequently cleared from circulation by renal elimination or reversibly transferred across the blood-brain-barrier by the common carrier of large neutral amino acids (LNAA_transp_). This reversible plasma-to-brain transport is defined through the unidirectional blood-brain clearances of FDOPA (K_1_^D^; mL/g min) and 3-O-MFD (K_1_^M^; mL/g min), and the corresponding rate constants for clearance back to circulation (k_2_^D^, k_2_^M^; min^-1^). FDOPA in brain tissue can be O-methylated at apparent rate constant k_5_^D^ (min^-1^) by COMT or decarboxylated at the rate constant k_3_^D^ (min^-1^) to form FDA. FDA is reversibly sequestered in vesicles by the vesicular monoamine transporter (VMAT) and then released into the synaptic cleft as part of both tonic and phasic dopamine release. FDA can then be reabsorbed into the presynaptic terminal and possibly be restored or metabolized by monoamine oxidase (MAO). Cytosolic FDA can also diffuse away from its source neuron to undergo metabolic destruction at another site; or it can be decomposed by MAO at rate constant k_7_^DA^ (min^-1^), yielding the acid metabolites [^18^F]fluorodihydroxyphenylacetic acid (FDOPAC) and [^18^F]fluorohomovanillic acid (FHVA). The acidic metabolites of FDA are together eliminated from brain by passive diffusion at rate constant k_9_^ACIDS^ (min^-1^). **B) Modelling [^18^F]fluorodopa (FDOPA) and 3-O-methyl-[^18^F]FDopa (3-O-MFD) kinetics.** FDOPA PET signal in striatum might require 3 different levels of modelling: a compartmental model representing FDOPA accumulation in brain (red compartments), a compartmental model representing the exchange between blood and brain of the FDOPA metabolite 3-O-MFD (blue compartments), and a compartmental model representing the clearance of [^18^F]fluorodopamine (FDA) and its acidic metabolites FDOPAC and FHVA (green compartments). Depending on duration of scanning or administration of peripheral COMT and DDC blockers prior to the PET scanning a combination of these models is used. None of these models include the coefficient of brain tissue methylation of FDopa k_5_^D^, because it is assumed to be negligible throughout the brain. FDA and its acidic metabolites can also occupy the same compartment and a common rate constant for their clearance from brain can be defined as k_clearance_ or k_loss_.

The use of FDOPA PET in discriminating response would require further validation on larger clinical datasets, aiming to support future individualized treatment and patient stratification across the different brain diseases. However, to reach clinical translation, FDOPA PET requires a suitable data infrastructure and robust analytical protocols to ensure high quality of the data, accurate quantification, and replicable results. As for any modern neuroimaging biomarker, the inhability to provide sufficient companion data and the lack of an analytical framework would hamper FDOPA PET applicability (Organization for Economic Cooperation and Development (OECD) 2011)(McAteer et al. 2021). An additional obstacle in the creation of such infrastructure is the inconsistency and variety in neuroimaging data format, which has a direct impact on the quality and confidence of the data. The creation of large neuroimaging repositories that gather data from multiple sites and sources inevitably faces the problem of data harmonization (Smith and Nichols 2018), and with the current fragmentation, it is very difficult (if not impossible) to create a single data management and analysis system that works for all the possible scenarios.

The reproducibility of the data analysis is another requirement for any clinical translation of a neuroimaging biomarker to be effective. Poor scientific reproducibility is in fact embedded in the complexity of the data as well as in their analysis pipelines, making difficult to guarantee transparent and certified analytical processes (Stupple, Singerman, and Celi 2019). For example, it is well-known that the neuroimaging results can be highly dependent on the analytical method chosen. A recent study, in which 70 independent teams were asked to analyse the same MRI dataset, led to significant discrepancies between execution and results, and demonstrates how the flexibility of the analytical approaches leads to important differences in the quantification of the data (Botvinik-Nezer et al. 2020). Similar findings were also observed for PET neuroimaging (Veronese, Rizzo, et al. 2021). In the case of FDOPA PET, there are several different analytical methods available, and ensuring reproducibility between these methods is far from trivial (Kumakura and Cumming 2009). Kinetic modelling and imaging pre-processing methods are necessary steps to isolate the biological components of interest from the measured FDOPA PET signal (**Figure 1B**). Furthermore, quantification of FDOPA PET imaging consists of the measurement of the activity of aromatic amino acid decarboxylase for dopamine production, which returns information about the functional integrity of the presynaptic dopaminergic synthesis (Kumakura and Cumming 2009). This information of interest needs to be isolated from the total measured PET radioactivity, removing the contribution of the tracer metabolism and non-specific binding (Kumakura and Cumming 2009). It follows that the statistical proprieties of the radioligand in term of reproducibility and biological variability are not sufficient to guarantee the applicability of the method in clinical setting, and both the tracer kinetic modelling and the imaging data analysis pipeline need to be validated as constitutive parts of the FDOPA PET imaging biomarker.

In this study we present and validate a new infrastructure for FDOPA PET neuroimaging. The project takes advantage of a large FDOPA PET data repository available at the Institute of Psychiatry Psychology and Neuroscience (IoPPN) at King’s College London. The first aim of this study is to harmonise the FDOPA PET data collected across several research studies. The second aim is to implement an automatic FDOPA PET data analysis pipeline, directly embedded in the data infrastructure, and to test its replicability and reproducibility, as well as its sensitivity to experimental and demographic covariates.

## METHODS

### FDOPA PET data acquisition

All FDOPA PET imaging sessions in the database were acquired with a continuous dynamic acquisition (no blood sampling), with scanning beginning with the tracer injection and lasting for 90-95 minutes. During this time, the participant was required to lie still in the PET scanner, with head rests to limit subject head motion. All participants received carbidopa (150 mg) and entacapone (400 mg) orally ~1 hour before imaging. Both drugs are used to increase the signal-to-noise (SNR) of the tracer uptake in brain tissue by reducing the peripheral formation of radiolabelled dopamine and the 3-O-methyl-[18F]fluorodopa brain-penetrating metabolite, respectively (Veronese, Santangelo, et al. 2021). The FDOPA tracer (injected dose ranging from 86.4 to 414.4 MBq,) was administered by intravenous bolus injection after the acquisition of a brain CT or MRI for attenuation correction, depending on the scanner availability at each imaging site. PET data reconstruction varied across imaging sites and scanner types, but all included correction for random noise, scatter and tissue attenuation.

### Data management infrastructure

Our data management infrastructure was built using XNAT imaging technology (Marcus et al. 2007). A bespoken installation of XNAT platform was deployed using the Neuroimaging Analysis Network at the Centre for Neuroimaging Sciences (King’s College London) to store for each subject’s demographic, clinical information and FDOPA PET imaging data. Representational State Transfer (REST) Application-Program Interface (API) was used to upload data.

For each subject, a scan imaging session was defined by a minimum amount of data which included dynamic FDOPA PET images (both attenuation-corrected and not attenuation-corrected), together with an ancillary file containing information regarding radiochemistry, date and time of scanning, timing of the acquisition and dynamic framing. Prior to storage in XNAT, the data were anonymized, harmonized in neurological convention, and corrected for radioisotope decay to ensure consistency across the data (**Figure 2**).

**Figure 2:**
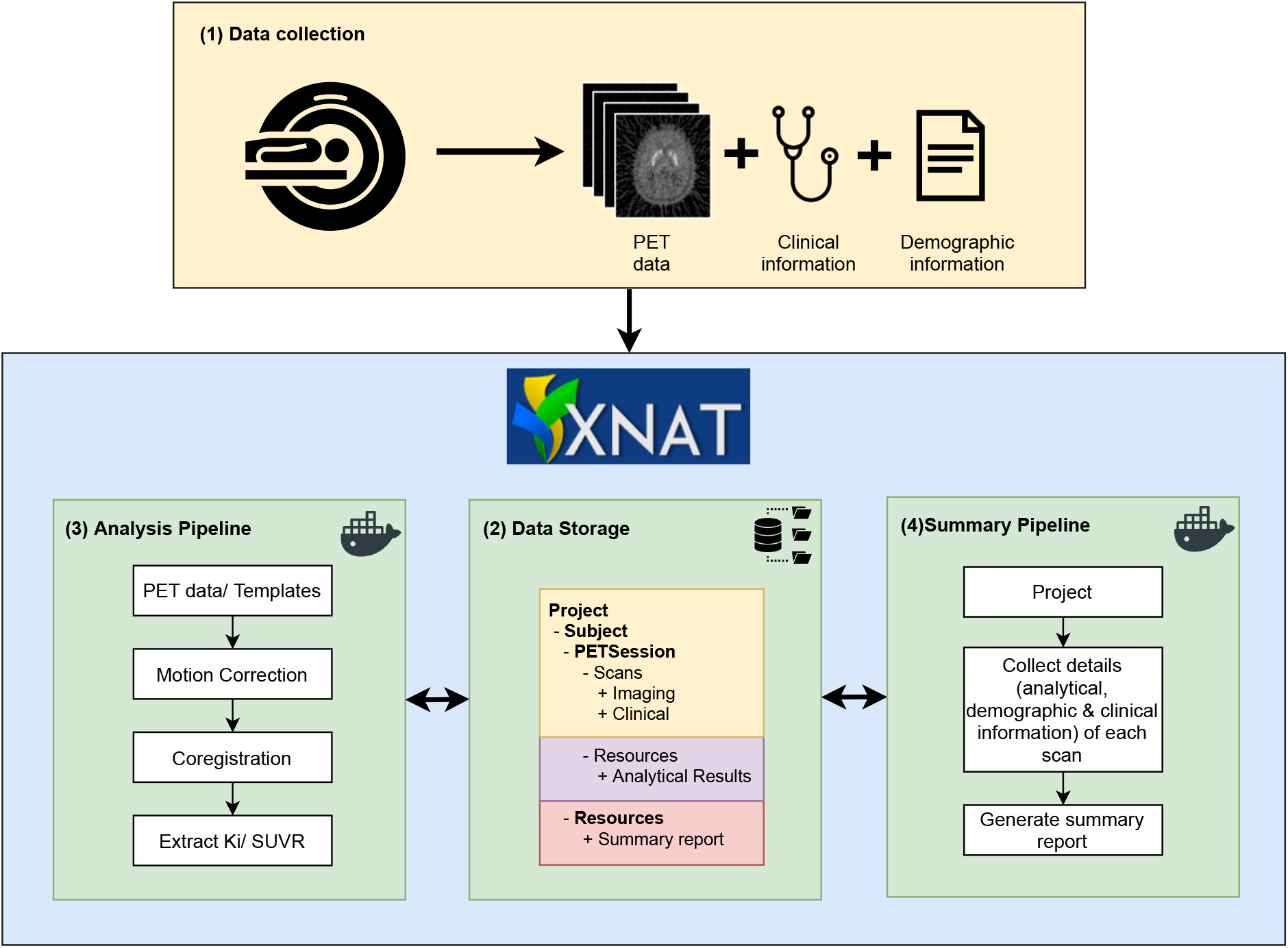
Schematic flowchart of the proposed infrastructure: the data are collected (1) and stored in XNAT (2). The main analysis pipeline for FDOPA PET quantification (3) and the summary pipeline (4) run automatically on the data, and the output are stored following the same data structure.

XNAT includes the Container Service plugin, which permits processes to be run on the stored data via REST API, using the processing utilities available in Docker containers (Merkel 2014). Taking advantage of this feature, analytical pipelines can be integrated in XNAT, allowing their automatic execution directly on the stored data and the storage of the outcomes at scan level (**Figure 2**).

### Automated analysis pipeline

The analysis pipeline was developed to align with our previous FDOPA imaging studies published by the Psychiatric Imaging Group (King’s College London) over the last two decades (Egerton et al. 2010, 2013, 2021; Howes et al. 2009, 2011, 2013; Jauhar, Veronese, Nour, Rogdaki, Hathway, Natesan, et al. 2019). In all these studies, the analysis pipeline was executed in MATLAB (Mathworks) and organised using a set of in-house scripts, which were manually executed for each individual scan.

The overall process of FDOPA PET data analysis can be described as follows. First, dynamically non-attenuated and attenuated FDOPA PET images are inputted into the pipeline. The non-attenuated dynamic images are motion corrected frame-to-frame to a single reference frame with a linear transformation. This reference frame is chosen at 15 min, as it represents an optimum trade-off between signal-to-noise ratio and radiotracer activity across all the brain tissues (**Supplementary Figure 1**). The motion information is then applied to the attenuated dynamic images, and the realigned frames summed together to obtain the motion corrected dynamic images. The information extrapolated during the motion correction step is used to quality control the data. This includes the “total motion”, estimated from geometrical realignment (both translation and rotation) of the individual PET frames, and the “number of spikes”, calculated as number of between-frames realignments exceeding a predefined threshold (5 mm, corresponding to the minimal spatial resolution detectable with standard clinical PET scanners). If one or more spikes are detected, the scan is flagged with potential motion artifacts. A tracer-specific template and atlas defining the striatum and cerebellum are co-registered to the attenuated PET images, using Statistical Parametric Mapping 12. To segment both the basal ganglia (main area of activity for FDOPA PET) and whole brain, two different atlases are used to extract the radiotracer activity at region of interest level: 1) an inhouse Montreal Neurological Institute (MNI)-based atlas including the whole striatum and its limbic, associative, and sensorimotor functional subdivisions as defined by Martinez et al (Martinez et al. 2003); 2) the adult maximum probability brain atlas developed by Hammers et al (Hammers et al. 2003). The Gjedde-Patlak Graphical approach (Patlak and Blasberg 1985; Patlak, Blasberg, and Fenstermacher 1983), with the cerebellum as reference region, is applied region-wise and voxel-wise to quantify Ki^cer^ (unit 1/min), a kinetic parameter used as a proxy of dopamine synthesis capacity (Kumakura and Cumming 2009). Prior to calculation of the Ki^cer^ parametric images, the images are denoised using a Chambolle Total Variation (Chambolle 2004) method. The parametric image for each scan is finally normalised into MNI standard coordinates using the participant’s PET summation image to calculate the image transformation field (non linear transformation). The Standardized Uptake Value Ratio (SUVR) is also calculated as ratio of the tracer activity to that in the reference region (i.e. mean cerebellar FDOPA PET activity). The interval 60-75 min after the injection of the radiotracer is used for this analysis, since in this time window the Gjedde-Patlak plot is linear and used to derive Ki^cer^ (Kumakura and Cumming 2009; Veronese, Santangelo, et al. 2021).

Starting from the available MATLAB code, a fully automated version of this analysis pipeline was written in Python using its standard libraries (NumPy, SciPy, etc.), and integrated in XNAT. All the pre-processing and processing steps executed in the MATLAB code (motion correction, atlas coregistration, region-wise and voxel-wise FDOPA quantification, MNI normalization, SUVR quantification) were replicated in the automated pipeline in Python. All the historical FDOPA scans were automatically re-analysed using the XNAT-based pipeline. To drive summary statistics for the entire database, an additional pipeline was integrated with the system. The pipeline permits to extract demographic, clinical, and analytical information in a summary report associated to each scan, automatically created, and stored in XNAT.

### Validation of the automated data analysis pipeline for FDOPA PET quantification

Data from two different datasets were used to assess the consistency of the XNAT-based pipeline with the MATLAB-based historical results and to test its reproducibility.

*Dataset 1* comprised FDOPA PET test-retest imaging data from 7 healthy controls, dynamically acquired using an ECAT/EXACT3D: Siemens/CTI (Knoxville, Tennessee) PET tomograph (spatial resolution: 4.8 (0.2) mm; sensitivity: 69 cps/Bq/mL). Approximately 150 MBq of 18F-DOPA was administered by bolus intravenous injection 30 seconds after the start of the PET imaging. Data were acquired in emission mode for 95 minutes, for a total of 26 time-frames reconstructed using a 3-dimensional reprojection algorithm. Full details of the research protocol and subject inclusion criteria are reported in the original reference (Egerton et al. 2010).

*Dataset 2* comprised FDOPA PET imaging data from 7 patients with psychosis before and after placebo, dynamically acquired using a Siemens Hi-Rez Biograph 6 PET scanner (Siemens, Erlangen, Germany) in 3D mode. Approximately 150 MBq of 18F-DOPA was administered by bolus intravenous injection after acquiring a CT scan for attenuation correction. PET data were acquired in 32 frames of increasing duration over the 95 min scan (frame intervals: 8 × 15 s, 3 × 60 s, 5 × 120 s, 16 × 300 s).

An independent dataset *(Dataset 3)* of 521 scans was used to evaluate agreement between XNAT-based and the already available MATLAB-based results, using the same analysis settings. A linear mixed model implemented with Jamovi (Version 2.0) was used to evaluate the effect of the detected number of spikes during motion correction on the FDOPA quantification.

### Identification of experimental and demographical covariates for FDOPA PET imaging

Baseline scans (no pre-scan intervention) of healthy controls were selected from the database to study the effect of experimental and demographic covariates on the FDOPA PET quantification. The final sample included 115 scans acquired from 3 different PET tomographs (Siemens Biograph 6 Hi-Rez, Siemens Biograph 6TruePoint, ECAT/EXACT3D) with an injected radioactivity below 200 MBq and acquisition time of 95 minutes. The experimental variables included tomograph type and injected radioactivity, and the demographic variables included participant gender, age and weight at the time of scanning. A similar analysis was then repeated on a subsample (N=103 scans), where the data were acquired using the Biograph PET tomographs (Siemens Biograph 6 Hi-Rez, Siemens Biograph 6 TruePoint) only.

### Statistical methods

For statistical analysis, GraphPad Prism v9 for Mac (GraphPad Software, La Jolla, CA) and SPSS (Version 27) were used. The Ki^cer^ estimates obtained from the XNAT and MATLAB analysis pipelines were compared using the Bland-Altman plot with 95% limits of agreement (Bland and Altman 1999). Correlation and mean absolute percentage difference between the two pipelines were also calculated for the following six regions of interest (ROIs): whole striatum, right striatum, left striatum, whole sensorimotor subdivision, whole limbic subdivision, and whole associative subdivision. These ROIs are commonly the primary areas of analysis in PET imaging dopamine studies (Martinez et al. 2003; Tziortzi et al. 2014). For both XNAT and MATLAB pipelines, test-retest reliability was estimated with the Intraclass Correlation coefficient (ICC) using a 2-way mixed-model in SPSS (version 27, IBM®), while the within-subject variation was calculated as the absolute percentage test-retest difference (Egerton et al. 2010). A p-value < 0.05 was considered statistically significant.

To investigate the effect of demographic and experimental variables on dopamine measures, a univariate general linear model (GLM) analysis was run with the K_i_^cer^ and SUVR of the whole striatum as dependent variable and both experimental and demographic variables as covariates.

## RESULTS

### Database

The final infrastructure included 892 FDOPA PET scans from 23 different studies. Both primary and secondary data were organized following the same structure and naming convention, which facilitates data management and ensures homogeneity across the database. After removing commercials studies for which we did not have permission for data reuse, the infrastructure consisted of 792 FDOPA PET scans from 666 individuals (female 33.9%, healthy controls 29.1%) collected from four different imaging sites between 2004-2021 (**Table 1**). The mean age of the participants was 28.7 years (*range = 18-65, S.D. = 8.5*) with a mean weight of 75.9 kg (*range = 38-136, S.D. = 17.3*). All scans were acquired from five separate tomographs (*Siemens Biograph 6 Hi-Rez, Siemens Biograph 40 TruePoint, Siemens Biograph 6 TruePoint, ECAT/EXACT3D, GE SIGNA PET/MR*) with a mean injected radioactivity of 188.2 MBq (*range = 86.4-414.4, S.D. = 80.1*).

**Table 1:** Demographic, experimental and imaging site information of the data (excluding the commercial studies) stored in XNAT.

| Attributes |  |
| --- | --- |
| ***Imaging Sites*** |  |
| Invicro, London | 455 |
| Imperial IMANET PET centre, London | 190 |
| Wolfson Molecular Imaging Centre, Manchester | 33 |
| Seoul National University Bundang Hospital, Republic of Korea | 114 |
| ***Controls (%)*** | 195 (29.1) |
| ***Age year***, mean (min-max) | 28.7 (18 – 65) |
| ***Weight kg***, mean (min-max) | 75.9 (38 – 136) |
| ***Gender female (%)*** | 243 (33.9) |
| ***Tomograph*** |  |
| Hi-Rez Biograph 6 | 366 |
| Biograph 40 Truepoint | 113 |
| Biograph TruePoint 6 | 70 |
| ECAT/EXACT3D | 49 |
| SIGNA PET/MR | 19 |
| ***Injected Radioactivity MBq***, mean (min-max) | 188.2 (86.4 - 414.4) |

Analysis of motion correction statistics highlighted that 6% of the scans (N=46) had at least one motion spike, and 2.8% (N=22) two or more spikes. The 85% of the total scans were patients. The mean and standard deviation of the motion detected for the healthy controls and patients (10.9±6.6 mm and 13.8±10.9 mm, respectively) were significantly different using the Mann-Whitney test.

There was a significant effect of the spikes on the Ki^cer^ estimates (*F=8.84, p<0.001*). Post-hoc tests showed that the Ki^cer^ significantly decreases for scans with two or three spikes (*p*<0.001) (**Figure 3**, graph on the left). Similarly, the SUVR significantly decreases for scans with five spikes (*p*<0.001) (**Figure 3**, graph on the right).

**Figure 3:**
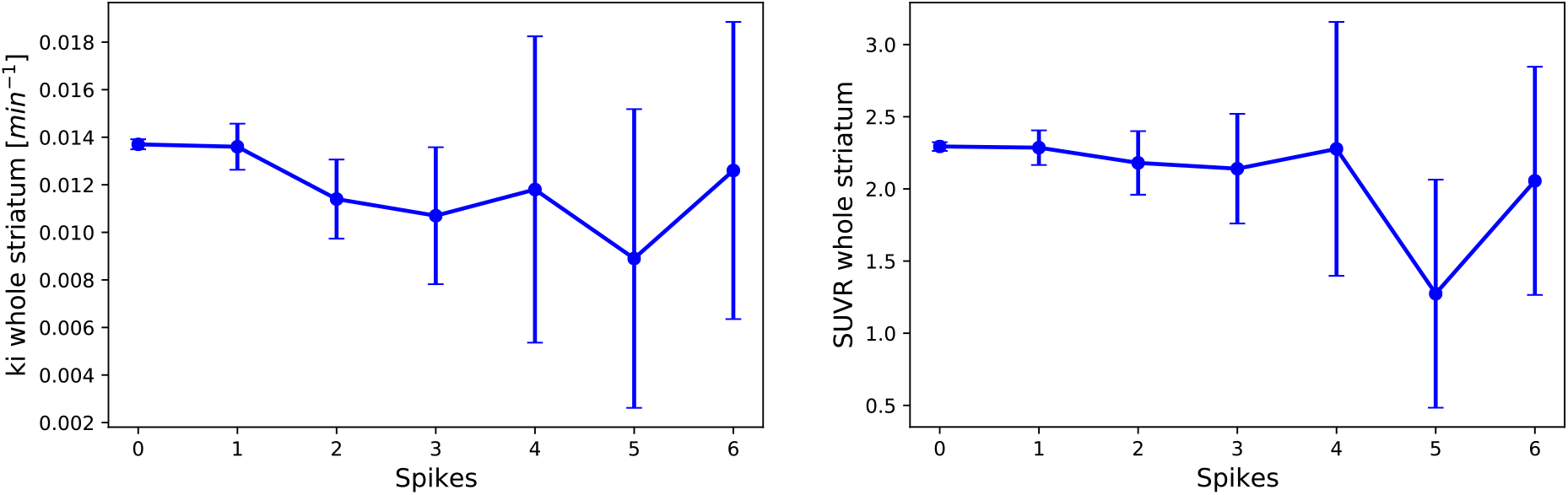
FDOPA quantification and spikes. Mean and 95% confidence intervals of the Ki^cer^ (A) and SUVR (B) of the whole striatum as a function of the number of spikes detected during the motion correction step of the analysis pipeline.

The Ki^cer^ distributions across the brain after removing scans with two or more motion spikes and/or visually-inspected corrupted tracer time-activity are presented in **Figure 4**. Among the ROIs, the highest estimates were reported in the whole striatum (*Ki^cer^ mean±SD: 0.0137±0.0015 min^-1^; min-max: 0.0102-0.0246 min^-1^ // SUVR mean±SD: 2.30±0.22; min-max: 1.71-4.11*). Outside the basal ganglia, the substantia nigra showed the highest signal (*Ki^cer^ mean±SD: 0.0072±0.0012 min^-1^; min-max: −0.0019-0.0122 min^-1^ // SUVR mean±SD: 1.53±0.15; min-max: 0.82-2.49*) followed by the pallidum, the amygdala, the thalamus and the prefrontal cortex. The occipital lobe, which is sometimes used as reference region for FDOPA PET quantification instead of the cerebellum (Dhawan et al. 2002), showed the lowest estimates in ratio to the cerebellum (*Ki^cer^ mean±SD: 0.0005±0.0004 min^-1^; min-max: −0.0009-0.0041 min^-1^ //SUVR mean±SD: 0.97±0.06; min-max: 0.79-1.72*).

**Figure 4:**
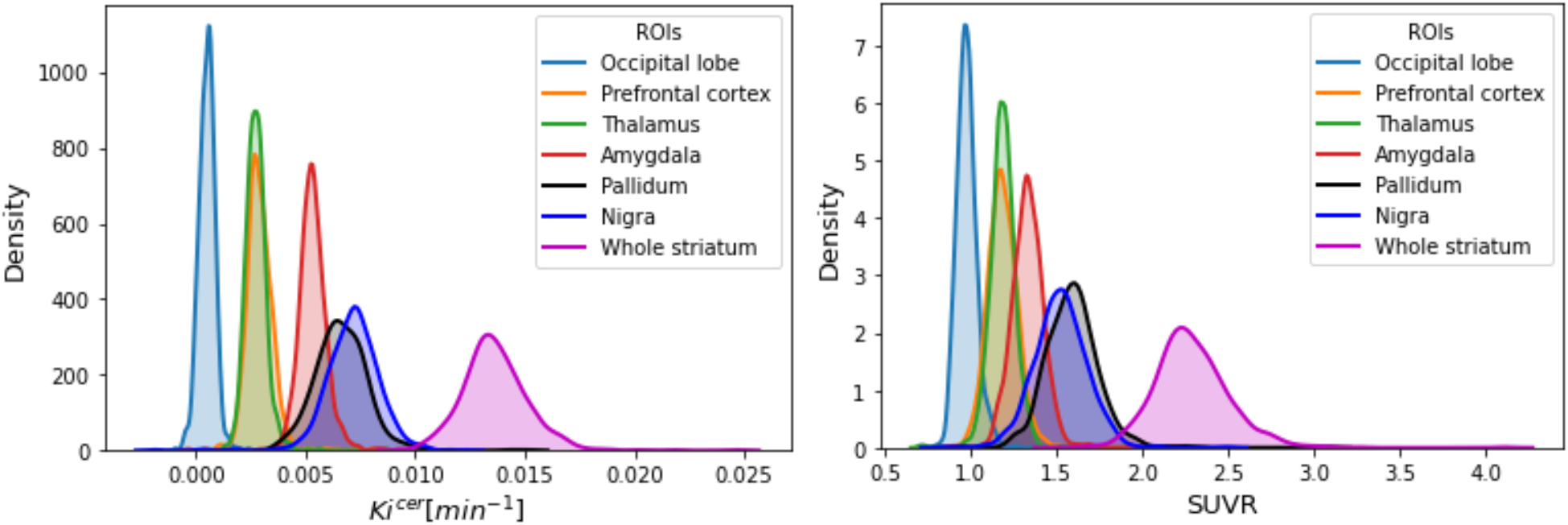
Distribution of Ki^cer^ and SUVR. Distribution of the Ki^cer^ (on the left) and SUVR (on the right) of the main region of interests (occipital lobe, prefrontal cortex, amygdala, substantia nigra, whole striatum) for all the subjects stored in XNAT.

### Validation of the automated data analysis pipeline for FDOPA PET quantification

The Bland-Altman plot confirmed good agreement between the XNAT and MATLAB pipelines for all the parameters, levels of analysis, datasets, and regions of interest. In the whole striatum, both XNAT-based Ki^cer^ and SUVR estimates are higher than the corresponding MATLAB ones within 5% mean relative difference (**Figure 5**). Consistent with the Bland-Altman analysis, XNAT-MATLAB Pearson’s correlation ranges from 0.64 to 0.99 for Ki^cer^, and from 0.79 to 1.00 for SUVR, with the lowest values for the the limbic subvidision and the highest for the whole striatum/associative subdivision (**Supplementary Table 1**). The mean absolute difference between the two pipelines ranges from 3.4% to 9.4% for Ki^cer^, and from 2.5% to 12.4% for SUVR (**Supplementary Table 1**).

**Figure 5:**
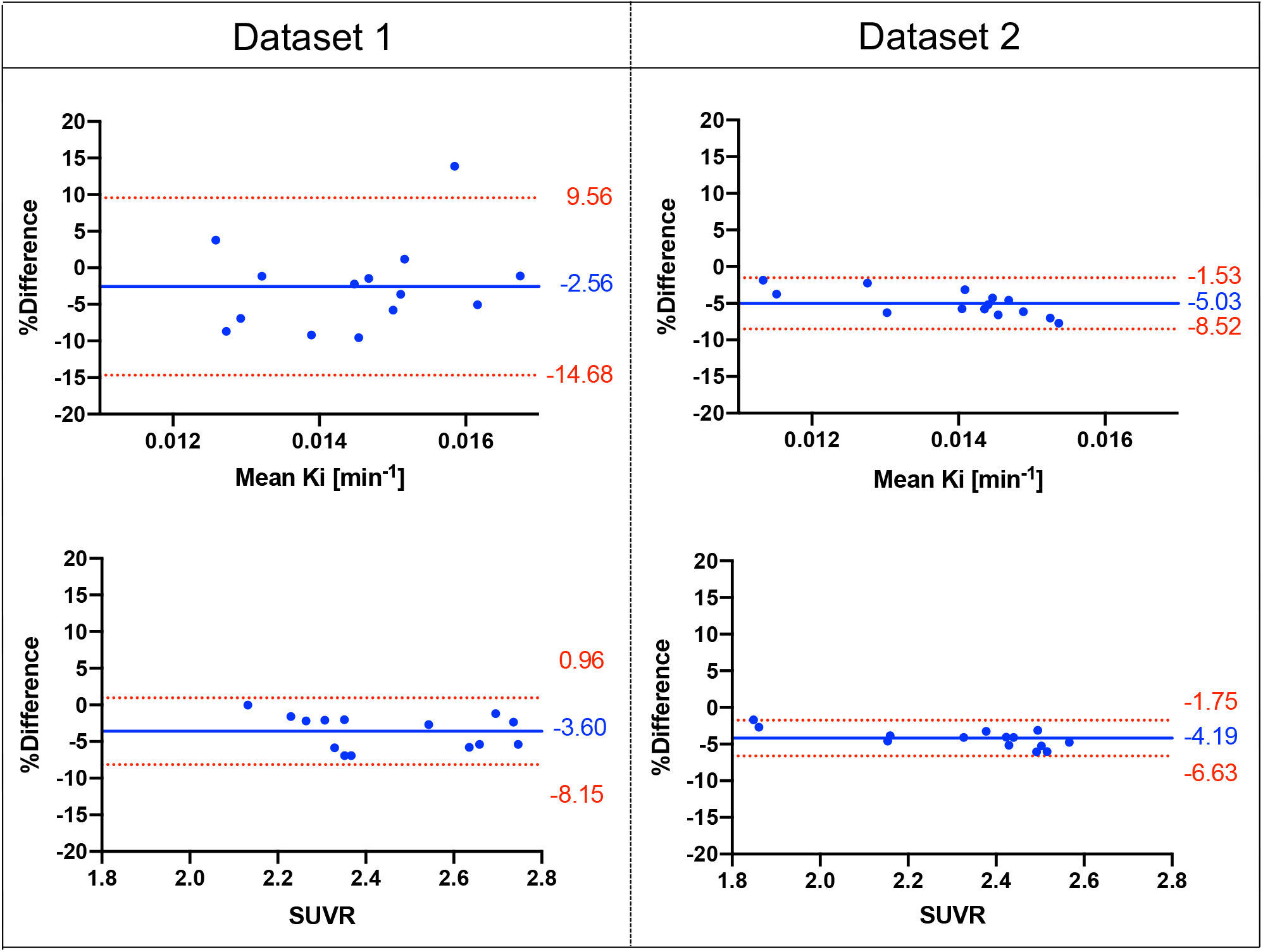
Bland-Altman plots comparing Ki^cer^ and SUVR estimates of the whole striatum with the XNAT-based and MATLAB-based pipelines, for both Dataset 1 and Dataset 2 [Difference (MATLAB – XNAT) vs. average]. The bias and 95% limits of agreement are reported in each graph.

In term of test-retest reliability and within-subject variation, the XNAT pipeline provided reproducibility and reliability of FDOPA PET in the striatum and its subdivisions (**Figure 6-7**) comparable to the ones previously described for non-attenuated FDOPA PET imaging pipelines (Egerton et al. 2010). By combining Dataset 1 together with Dataset 2 the ICC for the Ki^cer^ estimates ranged from 0.028 for the limbic subdivision to 0.929 for the right striatum. The %VAR for the Ki^cer^ estimates ranged from 3.7 for the right striatum to 19.3 for the whole limbic subdivision. For the SUVR, the ICC ranged from 0.79 for the limbic subdivision to 0.98 for the right striatum, while %VAR ranged from 2.5 for the right striatum to 6.0 for the whole limbic subdivision.

**Figure 6:**
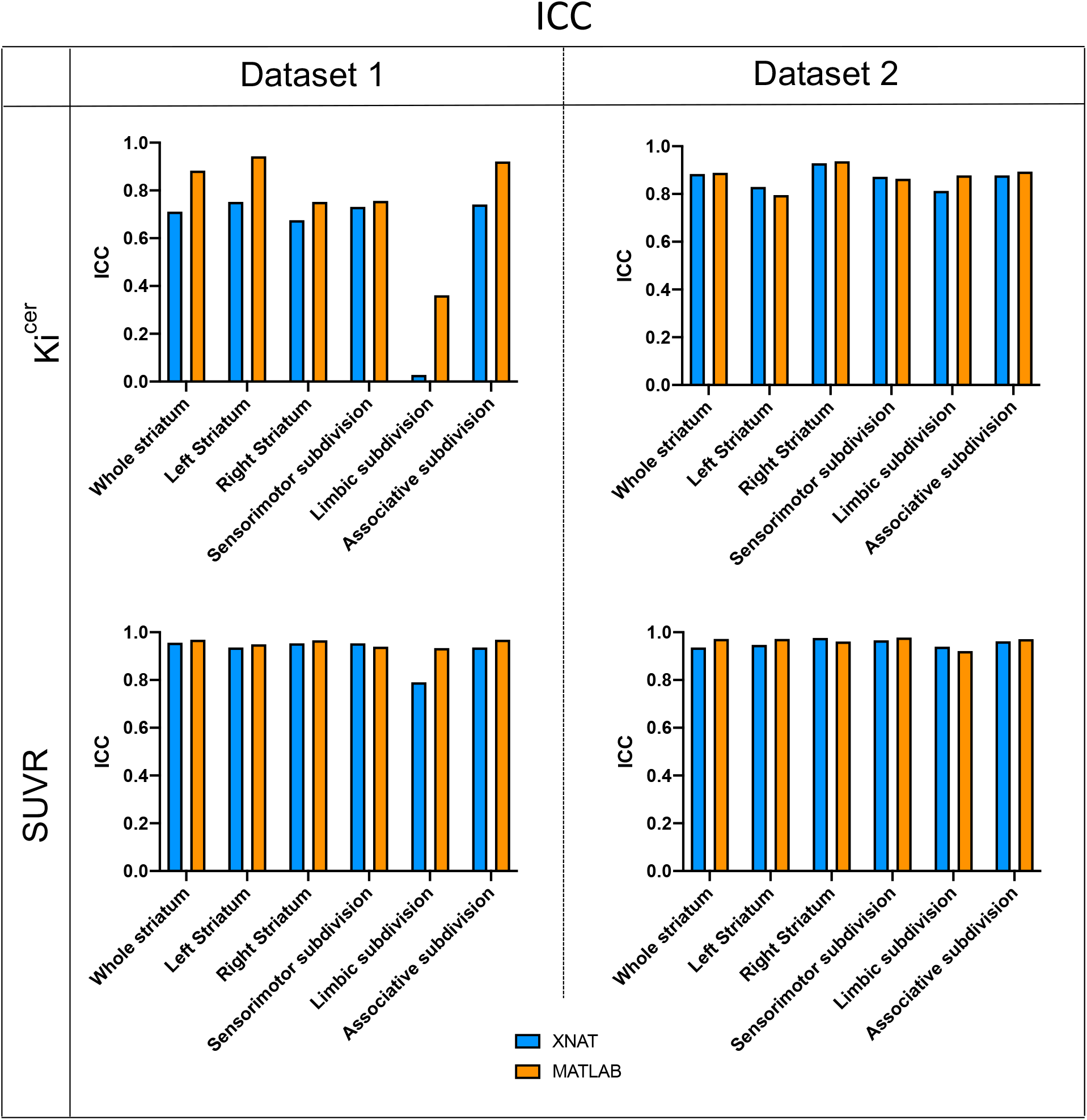
Comparison of test-retest characteristics shown as ICC between the XNAT (blue) and MATLAB (orange) pipelines for Ki^cer^ and SUVR of the whole striatum for both Dataset 1 and Dataset 2.

**Figure 7:**
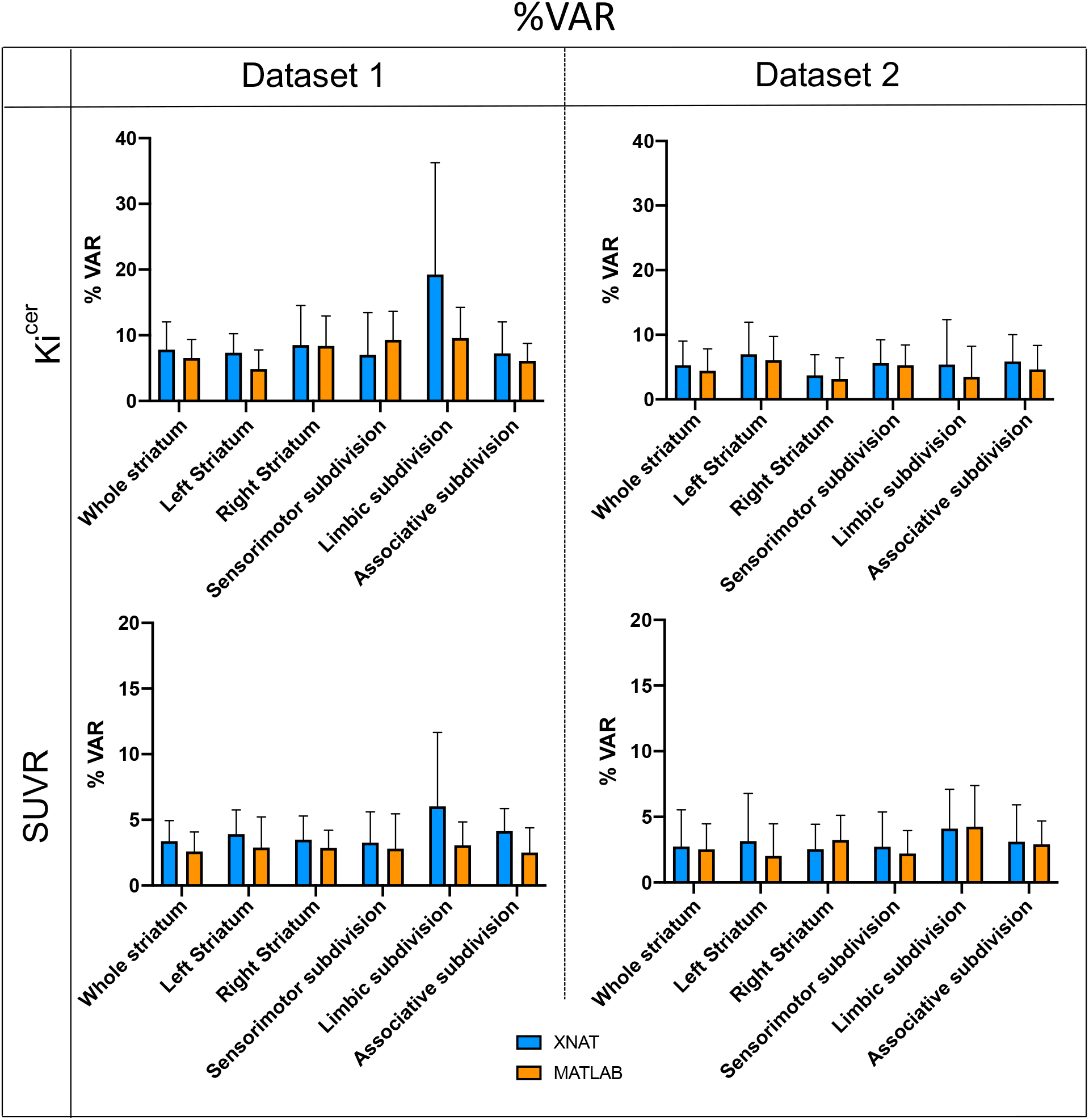
Comparison of test-retest characteristics shown as mean and 95% confidence intervals of %VAR between the XNAT (blue) and MATLAB (orange) pipelines for Ki^cer^ and SUVR of the whole striatum for both Dataset 1 and Dataset 2.

The consistency between the XNAT and MATLAB pipeline was further supported in an analysis of the independent and larger Dataset 3 (N=521), which showed a Ki^cer^ mean relative difference of 3.0±5.5% and a Pearson’s correlation of 0.86. About 71% of the scans reported Ki^cer^ relative differences less than 5%.

### Identification of demographical and experimental covariates for FDOPA PET imaging

There was a significant relationship between gender and striatal Ki^cer^ estimates (*F=10.7, df=1, p<0.001*), with dopamine synthesis capacity being higher in women than men (marginal means, women Ki^cer^: 0.0135±0.0015 min^-1^, men Ki^cer^: 0.0127±0.0013 min^-1^, effect size = 2.056E-5). Age, weight, injected radioactivity, and tomograph were not significantly associated with striatal Ki^cer^. A similar association between FDOPA PET and gender was also found when considering the subsample of scans (N=103) acquired only using the Siemens Biograph tomograph only (*F=10, p=0.002, female* Ki^cer^: 0.0135±0.0015 min^-1^, *male* Ki^cer^: 0.0127±0.0013 min^-1^) (**Table 2**). In contrast, none of the experimental and demographic variables were significantly associated with SUVR (**Table 2**).

**Table 2:**
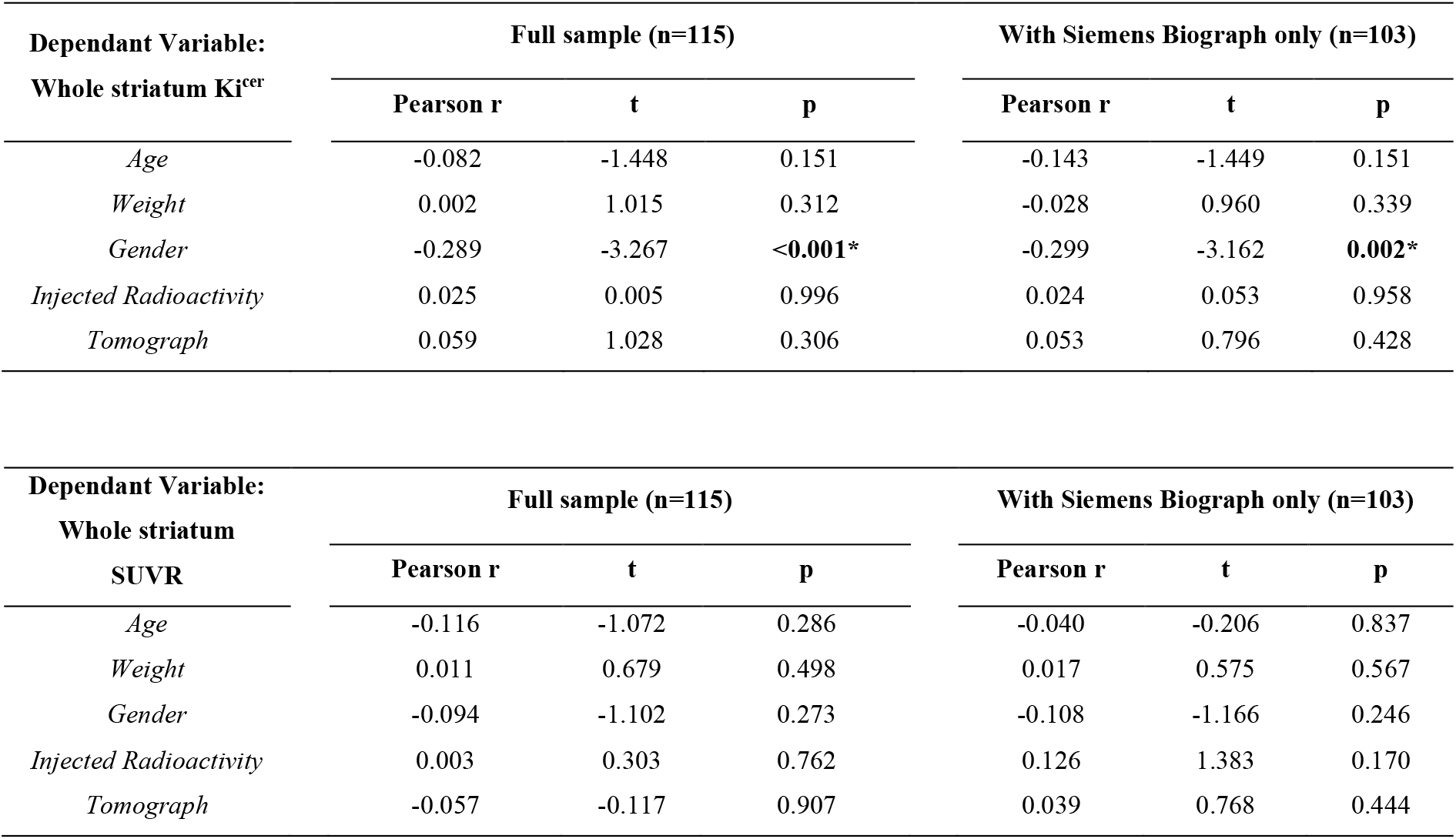
Association of demographical and experimental variables with Ki^cer^ and SUVR of whole striatum region.

| Dependant Variable:<br>Whole striatum Ki<sup>cer</sup> | Full sample (n=115) |  |  | With Siemens Biograph only (n=103) |  |  |
| --- | --- | --- | --- | --- | --- | --- |
|  | Pearson r | t | p | Pearson r | t | p |
| *Age* | -0.082 | -1.448 | 0.151 | -0.143 | -1.449 | 0.151 |
| *Weight* | 0.002 | 1.015 | 0.312 | -0.028 | 0.960 | 0.339 |
| *Gender* | -0.289 | -3.267 | **<0.001\*** | -0.299 | -3.162 | **0.002\*** |
| *Injected Radioactivity* | 0.025 | 0.005 | 0.996 | 0.024 | 0.053 | 0.958 |
| *Tomograph* | 0.059 | 1.028 | 0.306 | 0.053 | 0.796 | 0.428 |

| Dependant Variable:<br>Whole striatum<br>SUVR | Full sample (n=115) |  |  | With Siemens Biograph only (n=103) |  |  |
| --- | --- | --- | --- | --- | --- | --- |
|  | Pearson r | t | p | Pearson r | t | p |
| *Age* | -0.116 | -1.072 | 0.286 | -0.040 | -0.206 | 0.837 |
| *Weight* | 0.011 | 0.679 | 0.498 | 0.017 | 0.575 | 0.567 |
| *Gender* | -0.094 | -1.102 | 0.273 | -0.108 | -1.166 | 0.246 |
| *Injected Radioactivity* | 0.003 | 0.303 | 0.762 | 0.126 | 1.383 | 0.170 |
| *Tomograph* | -0.057 | -0.117 | 0.907 | 0.039 | 0.768 | 0.444 |

## DISCUSSION

In this study we characterize a standardised infrastructure for FDOPA PET neuroimaging to quantify brain dopamine function in living human brains. The platform was built from a harmonized FDOPA repository, integrating clinical and demographic information, together with FDOPA PET data. To our knowledge, this is the largest dataset of this type to date, including hundreds of scans with a very comparable acquisition protocol. The automated analysis framework for FDOPA PET quantification, directly embedded in the platform, ensured full control on the analytical process and replicable results on the stored data, confirming the good replicability and reproducibility of the FDOPA PET measures in the striatum and its subdivisions, expect for the limbic subdivision.

### Harmonization of data in big neuroimaging gathering

All the FDOPA PET imaging data were acquired using similar methods, which facilitated their integration in a unique storage infrastructure. However, the collection of FDOPA PET data from multicenter studies inevitably led to inconsistencies in the data structure and format used. To overcome these differences, the data were renamed using the same file name convention and manually converted to the same format (e.g. neurological convention, same image voxel size, data tracer decay corrected), prior uploading them in the XNAT platform. A rigid name convention was also used for the processed data obtained from the analysis pipeline, to ensure homogeneity across the entire dataset. This solution has been bespoken for the particular case study; however other name conventions might have been equally effective. The brain PET imaging community has lacked data format standards, and only very recently it has been recognised the need of more standardized data structure (Innis et al. 2007; Knudsen et al. 2020). The hope is that initiatives like PET-BIDS will address the heterogeneity of data organization by following the FAIR principles (findability, accessibility, interoperability, and reusability) (Nørgaard et al. 2021).

### Validation of the automated analysis framework for FDOPA PET quantification

The FDOPA Ki^cer^ and SUVR measures obtained with the XNAT pipeline have reproducibility and reliability in the striatum and its subdivisions comparable to the ones presented in Egerton et al in terms of test-retest variability and within-subject variation (Egerton et al. 2010), except for the limbic subdivision. The limbic subdivision showed in fact high Ki^cer^ variability and low ICC with the XNAT pipeline compared to the MATLAB for Dataset 1. These discrepancies in the Ki^cer^ quantification could be in part explained by the image pre-processing (motion correction, segmentation and coregistration). The limbic subdivision is a small region, more susceptible to motion artefacts and partial volume effects, and this can affect the amount of activity measured (Mawlawi et al. 2001). In addition, segmentation and coregistration in small regions can be challenging, expecially when using PET-based imaging data (Mawlawi et al. 2001). On the other hand, for the SUVR measures only the frames acquired in the interval 60-75 min after the injection of the radiotracer were considered, thus the effect of the image preprocessing could be less evident, explaining the improved ICC and variability in the limbic subdivision.

There was a good agreement in the FDOPA quantification in the automated compared to manual methods for both the test-retest datasets and the larger dataset. The percentage difference in the Ki^cer^ quantification between the manual and the automated method was within the acceptable threshold of 10% for 94% of the data. The percentage difference can be partially explained by the intrinsic complexity of the neuroimaging data analysis (Nørgaard et al. 2020). In this study, the XNAT-based pipeline was implemented from the available MATLAB code, but reproducing the exact results when using different analytical pipelines can be quite challenging due to the lack of standardized analytical pipelines and the lack of complete description of the used methodologies (Veronese, Rizzo, et al. 2021). The programming frameworks used, the computer environment and the choice of the pre-processing strategies are just few of the possible reasons behind different analytical outcomes. The discrepancies in the FDOPA quantification between the MATLAB and XNAT pipelines found in this study might come primarily from the pre-processing steps, which include motion correction, atlas coregistration and noise filtering. It is well-known that pre-processing steps are a critical part of a PET analysis framework, and small differences can impact the results (Nørgaard et al. 2019). In support of this aspect, we confirmed that the same Ki^cer^ and SUVR estimates were obtained when MATLAB-based preprocessed data were given in input to the XNAT-based pipeline (results not shown). It is also relevant to note that the two pipelines use different programming languages, MATLAB and Python, and different software packages, introducing another source of discrepancies in the data quantification. Unfortunately these are third-party components that are difficult to be controlled for. Hence the importance to keeping track of the software, libraries and packages used, as well as of all the steps used in the analysis framework, to ensure replicability and reproducibility.

### Technical and biological factors impacting FDOPA PET quantification

The availability of the proposed dataset allowed to characterise the distribution of the FDOPA PET signal across regions. The Ki^cer^ estimates in the whole striatum are the highest, ranging from 0.0102 to 0.0246 min^-1^ across individuals. In contrast to some recently published studies (Eisenberg et al. 2021; Sigvard et al. 2021) Ki^cer^ estimates smaller than 0.010 min^-1^ were only detected in scans with high head motion parameters, and rejected as outliers. Given the association between motion parameters and dopamine synthesis estimates, it becomes extremely important to physically limit head movements during the image acquisition. However, this can be uncomfortable and practically challenging. Alternative strategies to control and correct for data with high motion might be necessary, expecially in long acquisition. Usually the approach used to detect and exclude data affected by motion is study/site dependent and this can introduce further discrepancies in the results, which become hardly comparable. In addition, motion in patients and controls are different and this could have an effect in cross sectional studies.

The type of analysis pipeline is not the only factor that leads to FDOPA PET differences between published studies: in Eisenberg et al (Eisenberg et al. 2021), for example, the study participants did not received entacapone prior imaging acquisition, which is known to have a significant effect on FDOPA PET quantification both in animal (Léger et al. 1998) and humans (Keränen et al. 1993; Ruottinen et al. 1995). Differences in timing of acquisition (Cheng et al. 2020) are an additional source of variability to consider: kinetic parameters quantified by shorter PET acquisitions (e.g. 60 minutes) do not return the same estimates (Pan et al. 2005; Veronese, Santangelo, et al. 2021). Intermittent rather than continuous acquisition have also been proposed, but without a direct comparison with standard approaches the interpretation remains difficult (Cheng et al. 2020). Taken together, these factors highlight the importance of using a standardised acquisition protocol as well as common data analysis platform to compare results across studies.

The finding of a gender effect on the FDOPA measures, with higher Ki^cer^ and SUVR in female than men, is in agreement with established gender differences in brain dopaminergic activity (Laakso et al. 2002; Soutschek et al. 2017), where the number of participants used was relatively lower compared to the one used in the proposed study. Such differences might be explained by the effect of gonadal hormones, which modulate behavioral and neurochemical indices of activity in the striatum. Recent rodents studies have shown that, in female rats, estrogen increases presynaptic dopaminergic activity (Becker 1999), and a higher density of the striatal dopamine transporter is found in male rats (Rivest, Falardeau, and Di Paolo 1995; Walker et al. 2000), which may explain some of the differences found in this study, if such findings translate to humans. Gender is an important biological variable for different mental disorders and having a more comprehensive understanding on how it affects dopamine function would be of particular interest for the future development of individualized treatment response algorithms (Zachry et al. 2021).

### Limitations

In this study the data were manually quality controlled by visually inspecting the raw FDOPA tracer time-activity course, and the Ki^cer^ estimates and motion parameters obtained from the FDOPA quantification pipeline. This type of analysis is vulnerable to inconsistencies due to between-operator differences. Automatic pipelines for data quality control tailored to investigate specific characteristics of the data collected would reduce such issues (Pontoriero et al. 2021), but validated solutions are still missing.

Data provenance, defined as the documentation of where the data comes from and the processes and methodology by which it was produced, is fundamental to ensure full reproducible experimental and analytical processes (Chirigati and Freire 2017). The pipeline for FDOPA PET quantification, embedded in the platform, will need to be supported by documentation of the whole analytical process to further support and ensure full reproducibility of the scientific results. In terms of harmonization, the data were renamed using the same file name convention and manually converted to the same format to guarantee homogeneity across the database. However, the future aim is to use a unique standardized data format and structure, such as PETBIDS, which would facilitate the integration of data from different sources and sites and ensure a more reliable data harmonization.

The analysis pipeline implemented in this work follows the pipeline described in the variety of FDOPA imaging studies published by the Psychiatric Imaging Group (King’s College London). The pipeline could be further improved by integrating information from structural data like T1w MRI, which were not yet available for all the FDOPA PET scans, which could be used to enhance the atlas coregistration step.

XNAT was chosen as the platform for the implementation of the proposed infrastructure and no other alternatives were considered. XNAT was established in 2006 and since then it has been extensively used by research groups to host and collect clinical and other data associated with the initial raw imaging studies, enabling a broad range of collaborative research (Herrick et al. 2016). The system can be easily extended to other applications and new XNAT instances can be tailored to the different neuroimaging biomarkers, and the corresponding analytical methods can be integrated as automatic pipelines. However, XNAT is strongly imaging-driven and struggles to capture the variety and complexity of non-imaging data that are often acquired in modern experimental medicine studies. This information is important and could be integrated with neuroimaging data to provide a deeper individual phenotyping. The integration of this information in a unique system would permit to create a patient-centric platform, moving towards precision medicine.

## CONCLUSIONS

This study describes the development and analysis of a large repository for FDOPA brain PET data, which was undertaken with full control of the analytical process and produced robust and reproducible results. This unique harmonized FDOPA PET repository aims to further validate the accuracy and reliability of the method on a larger dataset. In addition, the availability of such large FDOPA PET repository aims to boost the future development of precision medicine applications across brain disorders, by investigating experimental and demographic covariates capable of explaining significant variability of dopamine synthesis capacity as estimated from FDOPA PET imaging.

## Acknowledgments

This study was funded by Wellcome Trust Digital Award (no. 215747/Z/19/Z) and supported by the National Institute for Health Research Biomedical Research Centre at South London and Maudsley National Health Service Foundation Trust and King’s College London. The views expressed are those of the author(s) and not necessarily those of the NHS, the NIHR or the Department of Health.

MV is supported by MIUR, Italian Ministry for Education, under the initiatives “Departments of Excellence” (Law 232/2016) and by the National Institute for Health Research Biomedical Research Centre at South London and Maudsley National Health Service Foundation Trust and King’s College London.

## Disclosure

OH and MV hold a patent application for the use of dopamine imaging as a prognostic tool in mental health (WO2021111116). The other authors do not report any conflict of interest in relation to this article.

## Supplementary material

**Supplementary Table 1:**
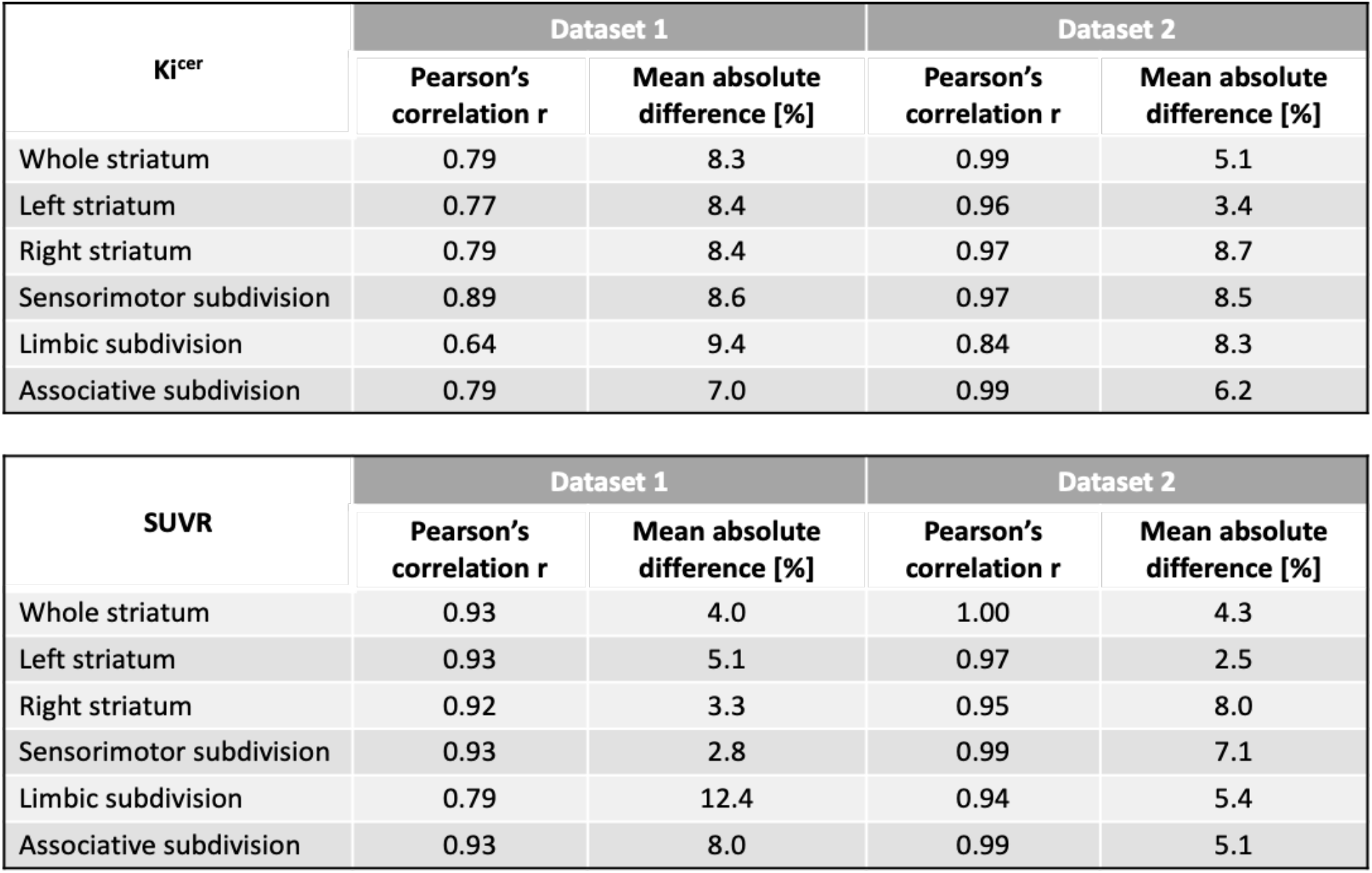
Correlation and mean absolute difference between XNAT and MATLAB pipelines for Dataset 1 and Dataset 2.

**Supplementary Figure 1:**
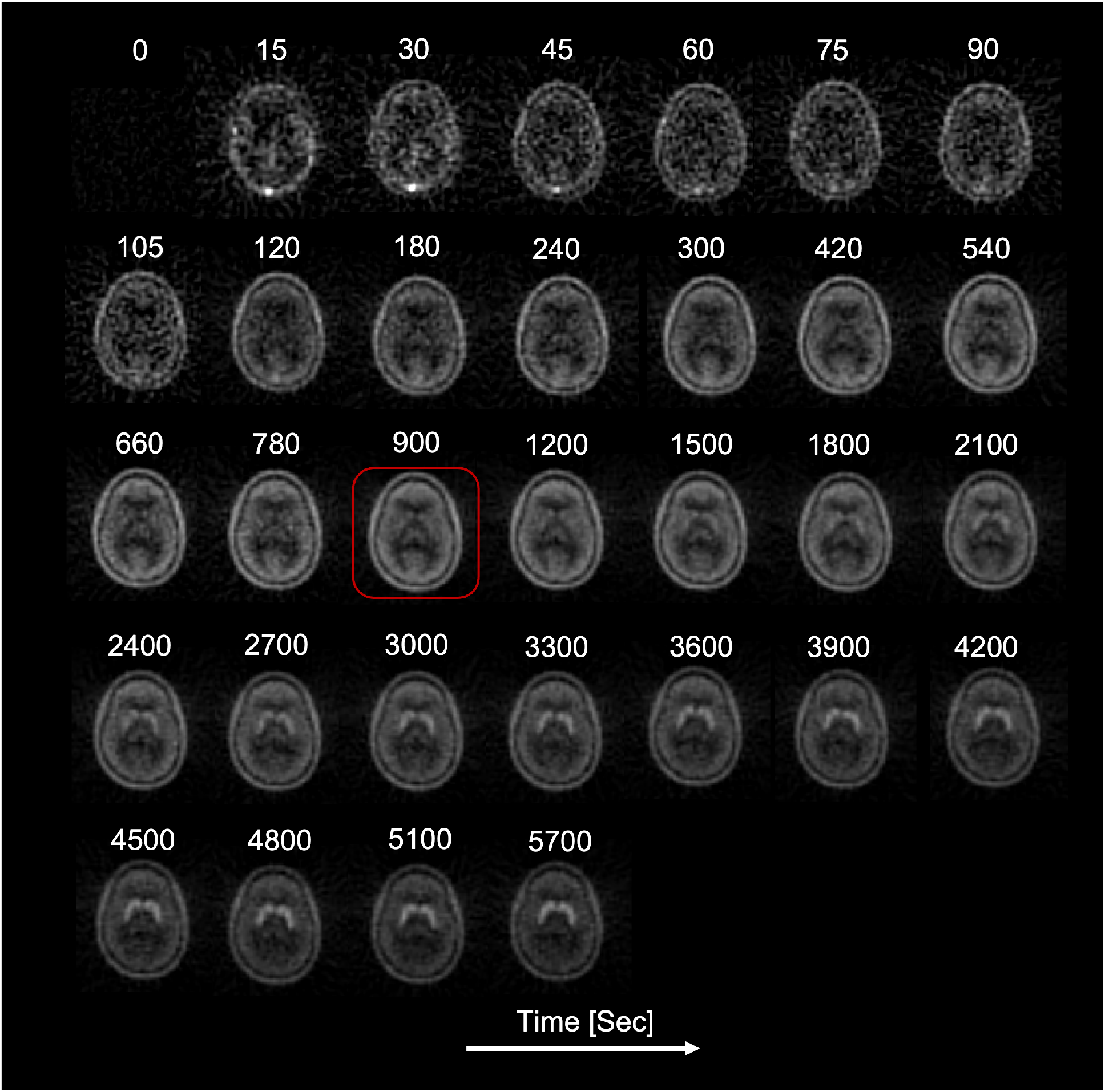
Single frame FDOPA PET images of a representative subject. The reference frame at 15 minutes used in the motion correction step is indicated in red.

## Notes

### Competing Interest Statement

The authors have declared no competing interest.

