## Supplementary figures and images for "Digital data repository and automatic analysis framework for FDOPA PET neuroimaging"

### Supplementary Figure 1

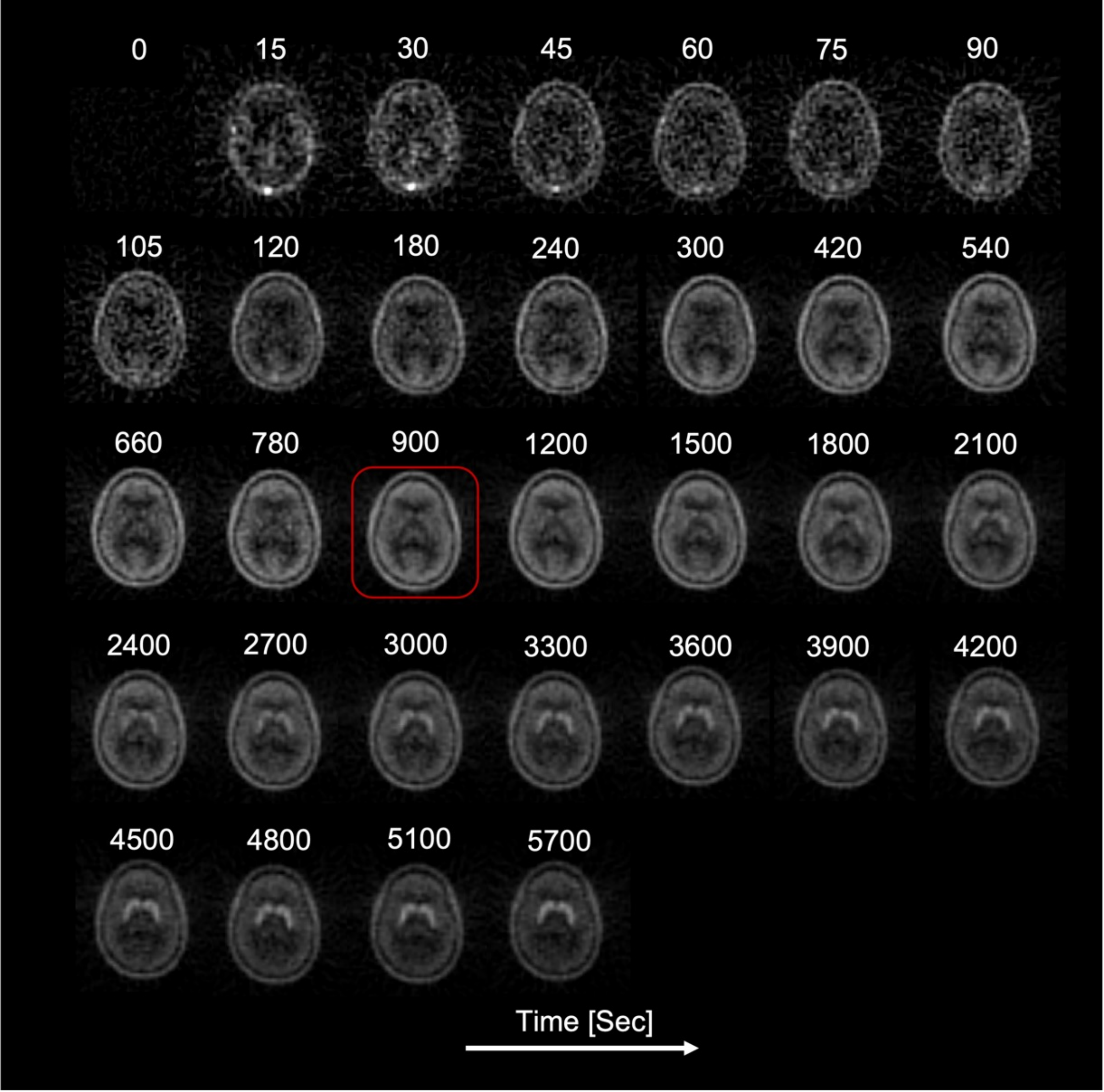
