## Supplementary Table 1 for "Digital data repository and automatic analysis framework for FDOPA PET neuroimaging"

| $Ki^{cer}$ | Dataset 1 | | Dataset 2 | |
|---|---|---|---|---|
| | Pearson's correlation r | Mean absolute difference [%] | Pearson's correlation r | Mean absolute difference [%] |
| Whole striatum | 0.79 | 8.3 | 0.99 | 5.1 |
| Left striatum | 0.77 | 8.4 | 0.96 | 3.4 |
| Right striatum | 0.79 | 8.4 | 0.97 | 8.7 |
| Sensorimotor subdivision | 0.89 | 8.6 | 0.97 | 8.5 |
| Limbic subdivision | 0.64 | 9.4 | 0.84 | 8.3 |
| Associative subdivision | 0.79 | 7.0 | 0.99 | 6.2 |

| SUVR | Dataset 1 | | Dataset 2 | |
|---|---|---|---|---|
| | Pearson's correlation r | Mean absolute difference [%] | Pearson's correlation r | Mean absolute difference [%] |
| Whole striatum | 0.93 | 4.0 | 1.00 | 4.3 |
| Left striatum | 0.93 | 5.1 | 0.97 | 2.5 |
| Right striatum | 0.92 | 3.3 | 0.95 | 8.0 |
| Sensorimotor subdivision | 0.93 | 2.8 | 0.99 | 7.1 |
| Limbic subdivision | 0.79 | 12.4 | 0.94 | 5.4 |
| Associative subdivision | 0.93 | 8.0 | 0.99 | 5.1 |
